# *E. coli*-produced nanobody fusions as a pan-serotype platform for AAV purification, quantification, and detection

**DOI:** 10.64898/2026.10.28.762627

**Authors:** Lucas Haverkamp, Marvin Fialho-Gerst, Kristian M. Müller

**Author notes:** Correspondence should be addressed to K.M.M.

## Abstract

Research, production and application of the recombinant adeno-associated virus (rAAV) require versatile tools for rapid and cost-effective advances. We developed fusion constructs with the pan-serotype, AAV-binding VHH-antibody-fragment named nanobody AAVX and devised procedures for AAV purification, quantification, and detection. Starting from an amino acid sequence, we cytosolically expressed in *E. coli* nanobody AAVX carrying various tags and fusion partners. Very efficient soluble production followed by a one-step immobilized metal-ion affinity chromatography (IMAC) yielded mostly >90% pure protein. Crosslinking AAVX with an agarose matrix enabled efficient purification of intact AAV2 and AAV5 capsids from cell lysates, as assessed by qPCR and ELISA. AAVX also supported effective AAV quantification by ELISA, as capture reagent and, when fused to luciferase, as detection reagent. Using tagged AAVX and streptavidin biosensors, we developed a rapid and label-free quantification assay of AAV capsids by biolayer interferometry (BLI). Finally, immunostaining with an AAVX-eGFP fusion visualized AAV capsids in fixed HEK-293 cells by fluorescence microscopy. Collectively, these results establish nanobody AAVX fusion proteins as versatile and readily customizable tools for AAV research and bioprocessing.

## Introduction

Adeno-associated viruses (AAVs) are established as potent viral vectors for gene therapy owing to their safety profile and high efficacy for *in vivo* delivery into various cell types.^1^ Currently, recombinant AAVs (rAAVs) are used in eleven FDA-approved products and in over 200 ongoing clinical trials.^2,3^ Despite this clinical success, widespread adoption is hampered by low production titers and AAV therapies face the challenge of immune responses, especially at higher doses.^4,5^ At the same time, rapid advances in the AAV field are driven by capsid engineering approaches aimed at modifying cell specificity, improving transduction efficiency, and modulating immune responses.^6–8^ These developments and the inherent challenges create a multifaceted research landscape that would benefit from broadly applicable but customizable analytical tools for AAV capsid detection, quantification, and localization.

The POROS CaptureSelect AAVX resin (Thermo Fisher Scientific) is used for affinity chromatography and is capable of capturing most natural and commonly used engineered capsids.^9,10^ Basis for the broad specificity is the variable domain fragment of a camelid heavy chain only antibody (VHH fragment or nanobody (Nb)), which binds the conserved DE- and HI-loops near the fivefold fold pore.^11^ The amino acid sequence of the nanobody AAVX was determined by Mietzsch *et al.* by nano-liquid chromatography tandem mass spectrometry after tryptic digest of the protein.^11^

Nanobodies are well established for diagnostic and therapeutic purposes and have proven to be a flexible research tool because of their small size, high affinity, and remarkable stability. Areas of application for nanobodies in research span detection in fluorescence microscopy, immunoassays, or biosensors, as well as characterization of proteins and protein interactions.^12–16^ Nanobody production is broadly established in diverse hosts including mammalian cells, plants, yeast, and prokaryotes.^17–20^ For research purposes, expression of Nb variants in *Escherichia coli* provides the benefit of easy and low-cost production with short turnaround times due to straightforward genetic modification and rapid growth.^21^ Most nanobodies carry at least one conserved disulfide bond, which can lead to improper folding and aggregation in insoluble inclusion bodies under standard expression conditions in the cytoplasm.^22,23^ The oxidizing environment of the *E. coli* periplasm creates suitable conditions for disulfide bond formation, but spatial constraints, translocation limitations, and insufficiency of periplasmic chaperones lead to compromised yields.^20^ Cytoplasmic expression of soluble nanobodies relies on disulfide bonds, which can be facilitated by engineering the cytoplasm to be more oxidative by either deletion of reductases as in commercially available strains like SHuffle and Origami, or expression of oxidases that catalyze disulfide bond formation like sulfhydryl oxidase (Erv1p).^24–26^ As we demonstrated previously, disulfide containing but very stable proteins may readily fold in the cytosol and express very well in standard *E. coli* strains. ^27,28^

Here, we describe an accessible recombinant production method for the nanobody AAVX and showcase some of its uses for flexible capsid analysis and purification. We immobilized the nanobody covalently on a crosslinked agarose-matrix via aldehyde-coupling to lysine residues. Furthermore, the nanobody was applied for AAV quantification as a ligand in biolayer interferometry (BLI) and in enzyme linked immunosorbent assay (ELISA).^29^ For the latter, the nanobody was used as a coating agent and in fusion with an enzyme based on the NanoLuc luciferase (nLuc) for detection.^30^ The NanoLuc has been shown to be a stable fusion partner and a highly sensitive reporter enzyme in ELISA.^31,32^ This approach was also combined with an ELISA protocol, which we developed in a previous study that utilized recombinantly expressed, soluble domains of the AAV cell entry receptor (AAVR) as coating agents.^33^ Lastly, the nanobody was genetically fused to eGFP and used to stain fixed AAV-producing HEK-293 cells to showcase a one-step immunostaining procedure.

## Results

### The Nanobody AAVX is expressed predominantly as soluble protein in the cytoplasm of *E. coli*

The amino acid sequence for the Nb AAVX was derived from the PBD entry 9DC3 and was ordered as a gene synthesis with optimized codons for expression in *Escherichia coli.* Tags and proteins providing additional functionality for purification, immobilization and detection were added as C-terminal fusion partners rather than N-terminal, due to potential interference with the capsid binding because of the proximity to the CDR3 region. All Nb variants were cloned into a pET21a plasmid and expressed in the cytosol of *E. coli* Origami (DE3), which provides inducible expression of the T7 RNA polymerase and supports formation of disulfide bonds in the cytosol through mutations in the thioredoxin reductase (*trxB*) and glutathione reductase (*gor*). Various Nb AAVX fusion constructs were designed for different purposes, while all constructs carried the effector at the C-terminus of the nanobody and histidine tag as shown in Figure 1A. Constructs were produced at small scale in shake flasks at 24°C and purified by a one-step immobilized metal-ion affinity chromatography (IMAC). AAVX-His had no additions besides the His-tag with the purpose of being used for characterization of expression behavior in *E. coli* and as a coating agent in ELISA. AAVX-nLuc carried a NanoLuc luciferase for quantification *via* ELISA and AAVX-GFP had an added eGFP for detection and localization in fluorescence microscopy. The AAVX-dHis construct was initially designed for strong binding to Ni-NTA resin for chromatography purposes and was based on a study by Khan et al. that incorporated two hexahistidine tags separated by a Strep-tag I for superior affinity to the resin.^34^ Figure 1B shows that the exemplary Nb AAVX-NanoLuc-luciferase fusion protein (AAVX-nLuc) is overexpressed upon induction and that the majority of the protein remains soluble after cell lysis. All constructs behaved similarly during production and purification as depicted in the supplementary materials (Figure S1). After Ni-NTA chromatography, Nb AAVX constructs were obtained at high purities (AAVX-His, - nLuc, -dHis > 90%, -GFP >80%), as determined by densitometry from the SDS-gel (Figure 1C), with yields of 18 mg per 1 OD_600_ and liter *E. coli* culture for the exemplary AAVX-nLuc. The purified Nb constructs showed good thermal stability with a T_m_ at 77.25 °C measured in a thermal shift assay (Figure S2).

**Figure 1:**
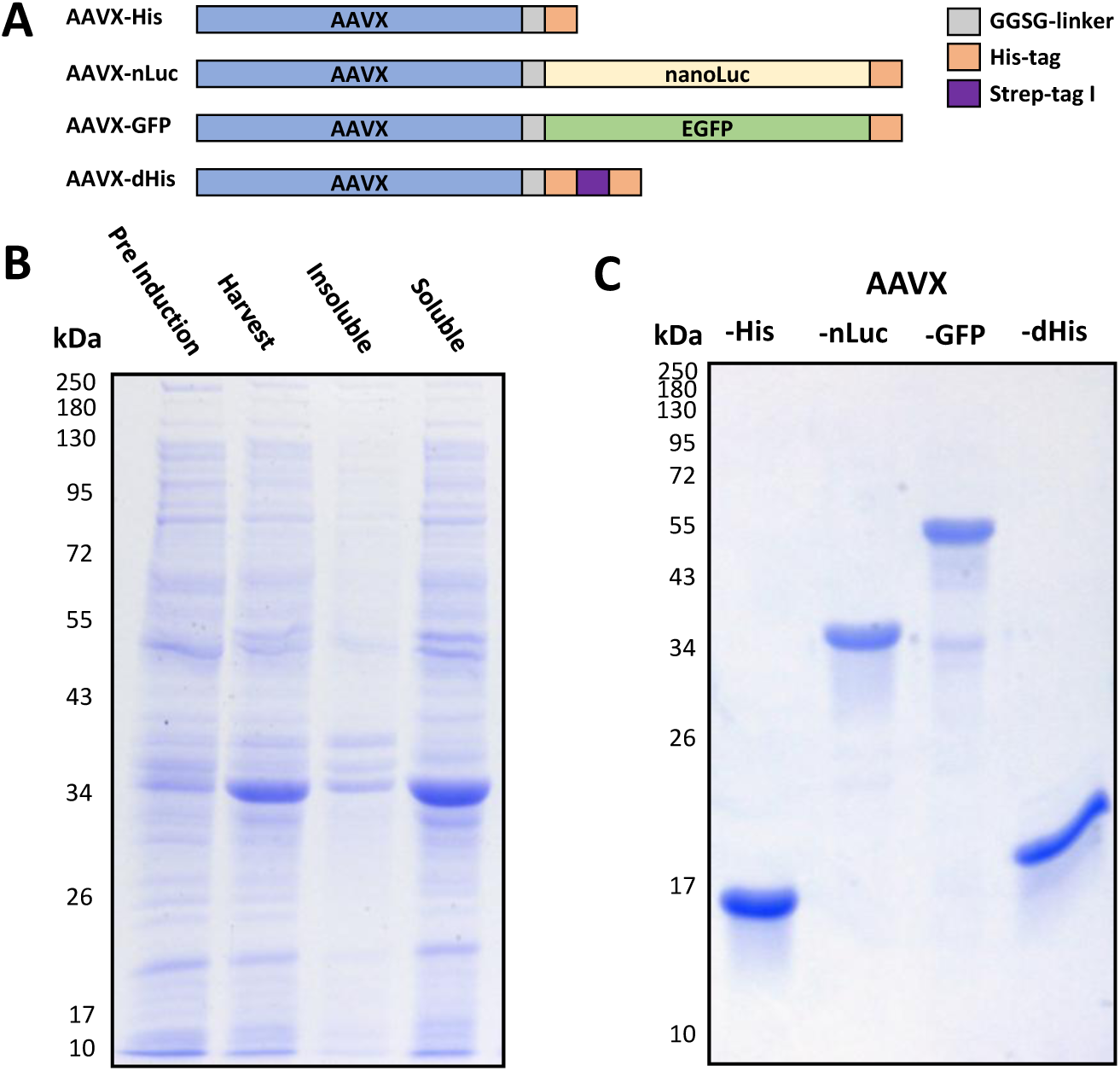
Expression and purification of Nb AAVX variants. (A) Scheme of Nb AAVX constructs with C-terminal fusions; - His: pentahistidine tag; -nLuc: NanoLuciferase; -GFP: eGFP; -dHis: dual hexahistidine tag interspersed by Strep-tag I. (B) SDS-gel displaying the production of AAVX-nLuc Fusion protein in E. coli Origami (DE3) with normalized samples; Pre Induction: Whole cell protein of E.coli before induction; Harvest: Whole cell protein of E.coli 18 h after induction; Insoluble: pelleted cell debris after cell lysis; Soluble: soluble proteins after cell lysis. (C) SDS-gel of Nb AAVX variants; 5 µg of IMAC-purified protein was loaded per lane.

### Nanobody constructs can be used for AAV affinity purification from cell lysate

To utilize the AAVX nanobody for purification, initially a non-covalent immobilization approach was investigated wherein the dual hexahistidine-tagged nanobody was bound to Ni-NTA agarose directly from cell lysate. This approach remained unsuccessful due to a lack of elution conditions, that efficiently eluted AAV particles without concomitant leaching of the nanobody (Figure S3). To circumvent these problems, an AAVX-GFP variant was immobilized by covalent coupling to a crosslinked agarose matrix with minimal loss during coupling and subsequent wash steps. AAV particles were efficiently bound by the resin by a batch chromatography and only a small fraction (approx. 0.1% of the eluted fraction) was lost during sample loading and wash steps (Figure 2A).

**Figure 2:**
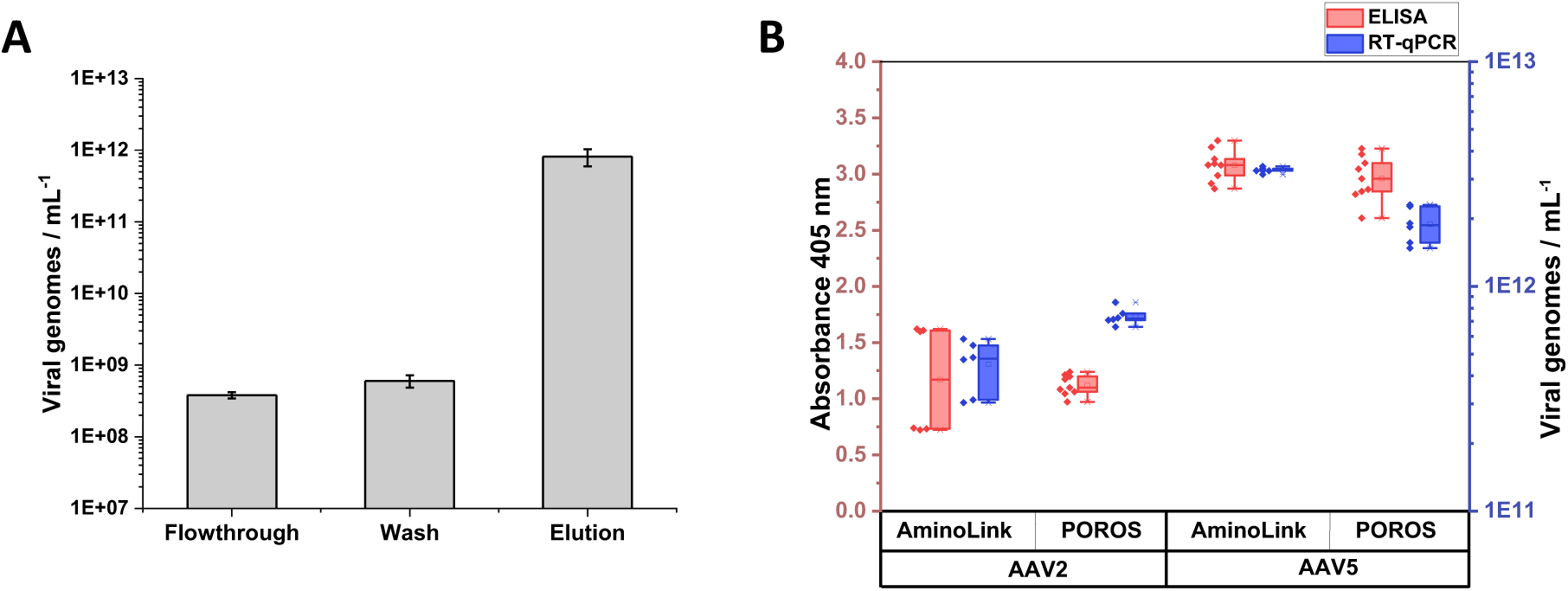
Purification of AAV particles with AAVX-AminoLink resin. (A) qPCR data of purification process of purified AAV5 particles (B) comparison of capsid ELISA signals and viral genome yields by qPCR after batch purification of rAAV from crude cell lysates with CaptureSelect AAVX affinity resin or AAVX-GFP immobilized on AminoLink resin (biological n=2 or n=3, technical n=3). A significance test was not performed to avoid overinterpretation.

For comparison with the commercial POROS CaptureSelect AAVX affinity resin, cell lysate samples of HEK-293 cells producing AAV2 or AAV5 were added to the resin for a batch purification. Small scale purification yielded similar results for each matrix material (Figure 2B). For AAV2, the commercial resin yielded slightly higher viral titers, whereas the AAVX-GFP performed better for AAV5 capsids. AAV2 as well as AAV5 samples did not differ significantly in capsid yield when comparing both matrices.

### Biolayer Interferometry measurements with the AAVX nanobody as ligand enable AAV quantification

BLI is a real-time, label-free, surface-based interaction analysis method, which allows for quantification when a calibration curve is available. Measurements of an association and dissociation phase provide affinity- and dissociation constants for a ligand-analyte pair. The AAVX-dHis construct also carries a Strep-tag I, which we used to immobilize the nanobody on streptavidin-coated biosensors (for details also see discussion section). AAV capsids added to the surface and bound by the nanobody lead to a proportional binding response depending on capsid concentration (Figure 3A).

**Figure 3:**
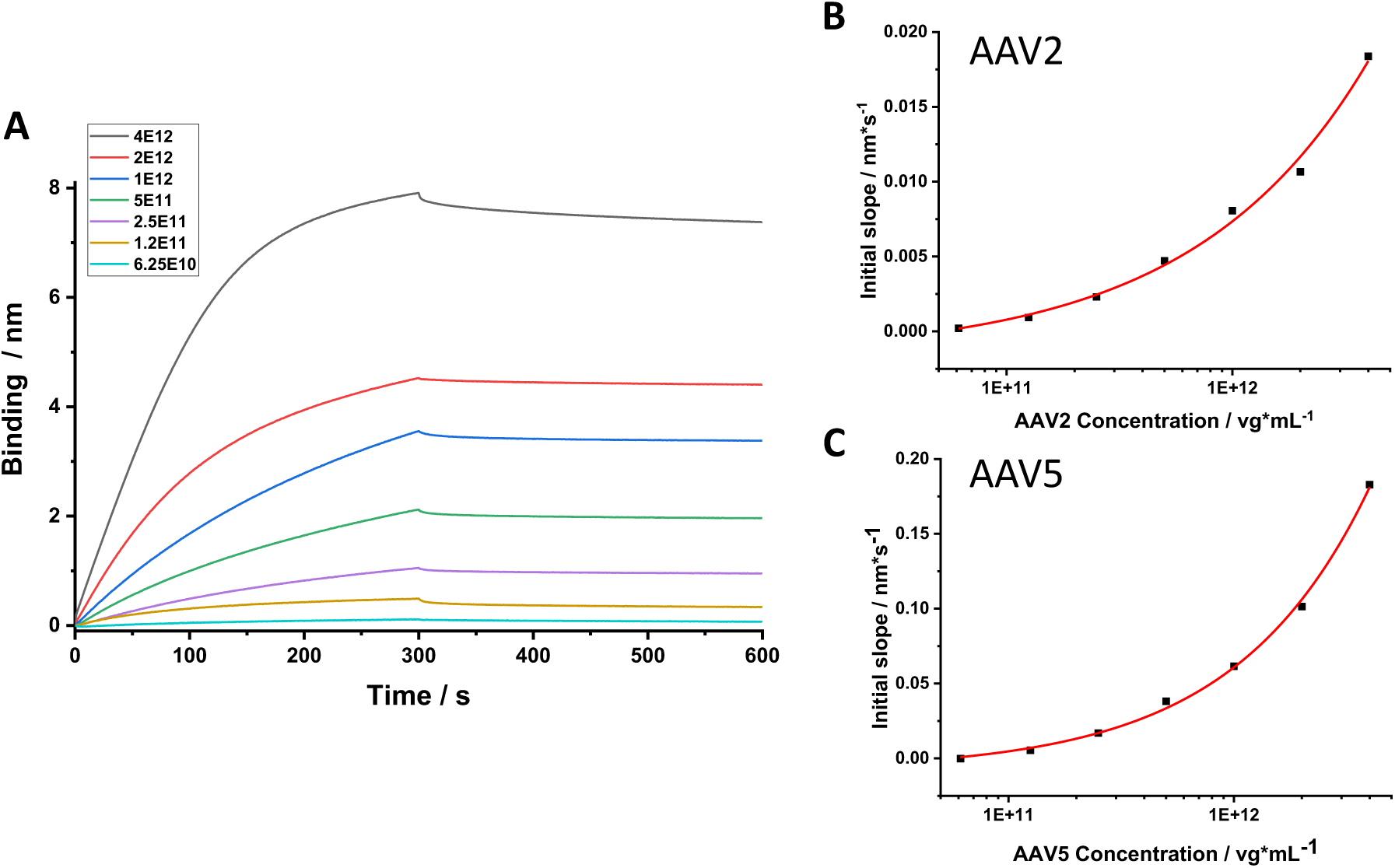
BLI measurement of Nb AAVX and AAV2 or AAV5 (A) sensorgrams of association and dissociation of AAV2 concentrations ranging from c.25×10^10^ to 4×10^12^ vg/mL (B) Initial slope of AAV2 association, fitted with a 4PL fit, R^2^ = 0.SS (C) Initial slope of AAV5 association, fitted with a 4PL fit, R^2^ = 0.S8.

To test whether calibration curves can be established, AAV2 and AAV5 capsids were applied in concentrations ranging from 6.125×10^10^ - 4×10^12^ vg/mL (1×10^-10^ mol/l – 6.6×10^-9^ mol/l). The slopes of the initial, linear binding response related to the apparent association constant (k_ass_) were extracted (Figure S5), plotted versus concentration and fitted with a 4 parameter-logarithmic fit to generate a standard curve for the quantification of unknown capsid concentrations (Figure 3B, C).

From the sensorgrams, next to the apparent association also the dissociation rate constants (k_diss_) were determined and the resulting apparent K_D_ for AAV2 that was in the picomolar range. The calculation of the apparent kinetic values did not factor in the multivalency of the AAV-AAVX-Streptavidin interaction and likely significantly overestimated the true affinity precluding a simple reporting.

### AAV quantification by ELISA with Nb AAVX for coating and/or detection

The Nb AAVX was utilized in sandwich ELISAs either for capturing by direct immobilization to the microtiter plate or for detection in a fusion variant with an enzyme based on the NanoLuc luciferase (nLuc). For capturing, Nb AAVX was adsorbed and AAV2 and AAV5 particles in concentrations ranging from 6.25×10^7^ to 4×10^9^ vg/mL were applied and subsequently detected with a primary, capsid specific mAb and secondary HRP-conjugated Ab. ELISA of AAV2 (Figure 4A) showed a better dynamic range across the samples, but lower absolute signal at higher concentrations in comparison to the AAV5 detection (Figure 4B).

**Figure 4:**
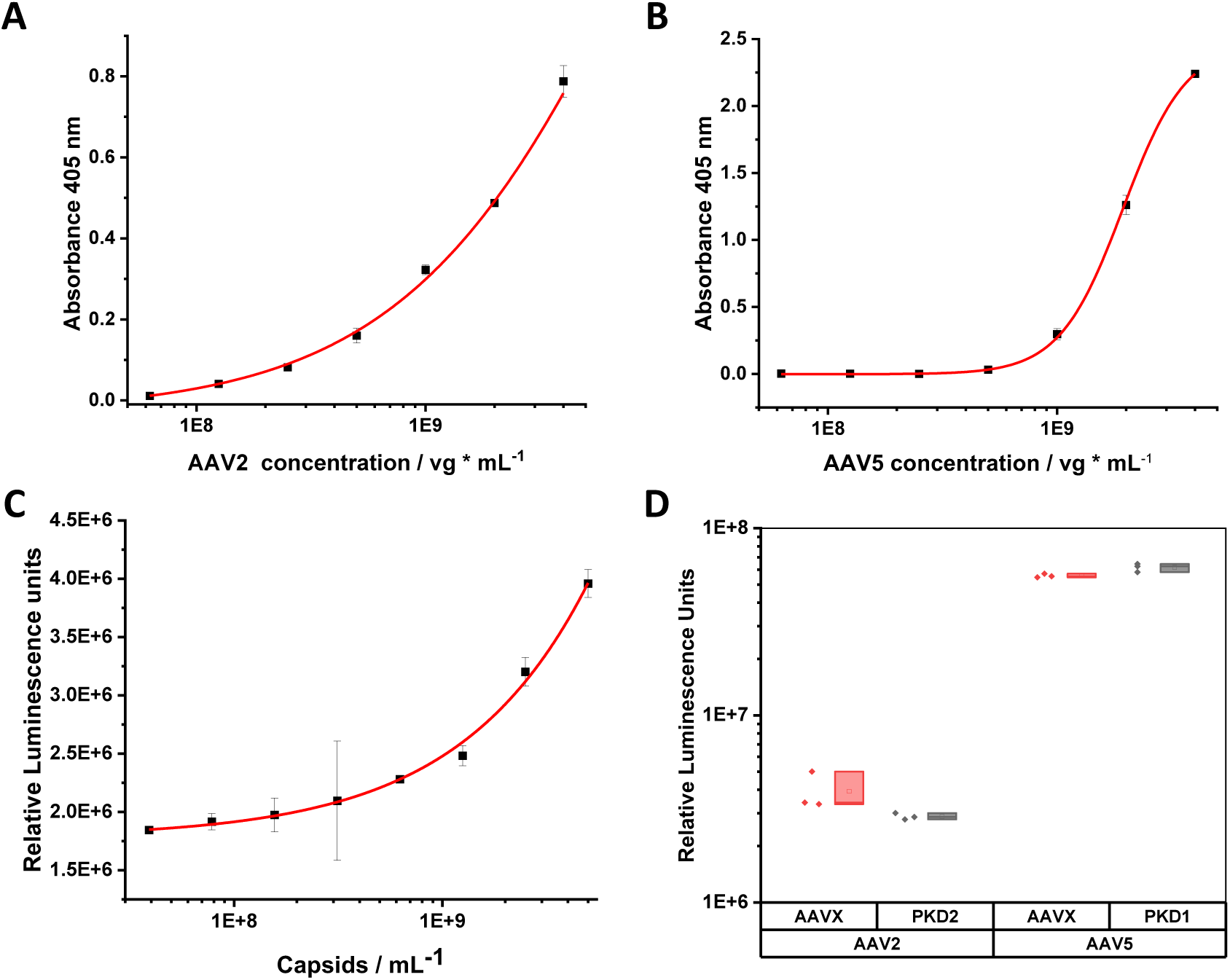
Sandwich ELISA of AAV capsids in Sc-well format. (A) Wells coated with 1 μg Nb AAVX, addition of AAV2 capsids and A20 as primary and anti-mouse HRP-conjugated as secondary for detection with ABTS, fitted with 4PL fit, R^2^=0.SS. (B) Wells coated with 1 μg Nb AAVX, addition of AAV5 capsids, primary mAb ADK5a, and secondary Ab anti-mouse HRP-conjugated for detection with ABTS, fitted with 4PL fit, R^2^=0.SS. (C) Wells coated with 1µg of AAVX, addition of empty AAV2 capsids from a standard (Progen) and detection with AAVX-nLuc, fitted with 4PL fit, R^2^=0.SS. (D) Wells coated with 1 μg Nb AAVX or PKD2-MBP or PKD1-MBP to capture AAV2 or AAV5 (2.5×10^S^vg/well) and detection with AAVX-nLuc. (n = 3).

For the evaluation of capturing and detection solely relying on the Nb construct, Nb AAVX was adsorbed, empty AAV2 capsids in a concentration range of 3.91×10^7^ to 5×10^9^ capsids/mL were applied and subsequently detected with Nb AAVX-nLuc. This sandwich ELISA with direct detection yielded a signal related to concentrations that was fitted with a 4-parameter logistic fit providing a calibration curve for determining capsid titers (Figure 4C).

Finally, we compared the sandwich ELISAs using Nb AAVX with our previously developed ELISA format, in which the AAV was captured by recombinantly expressed domains of the AAV-receptor.^33^ The PKD2 domain of AAVR binds to AAV2 and the PKD1 domain to AAV5 and both were produced in *E. coli* as maltose binding protein (MBP) fusion proteins and purified. In this ELISA, either Nb AAVX or PKD2 or PKD1, respectively, were coated and AAV2 or AAV 5, respectively, added and Nb AAVX-nLuc was used for detection in all cases (Figure 4D). Nb AAVX and the PKD domain capturing yielded comparable signals with PKD2 signal being only slightly below and PKD1 slightly above the Nb AAVX signal.

### AAV detection in cells by immunostaining with AAVX-GFP

Next, we tested whether the fusion protein of Nb AAVX with eGFP, named AAVX-GFP is suited to detect AAV particles in cells. HEK-293 cells were transfected with AAV2 pRepCap and pHelper, fixed 48 h post transfection, permeabilized, and stained with AAVX-GFP in concentrations of 10 µM or 50 µM. In fluorescence microscopy images, for both concentrations the GFP signal localized predominantly to DAPI-stained nuclei and occurred in clusters indicating expected AAV production in nucleoli. Staining intensities correlated with applied concentrations (Figure 5, last two lanes). To determine nonspecific staining at high concentrations, untransfected cells were stained with 50 µM AAVX-GFP and even at this high concentration no background was detected (Figure 5, first lane). As additional comparison, the commercial anti-AAV2 mAb A20 was used to stain transfected cells and the detection pattern was relatively similar to the AAVX-GFP staining (Figure 5, second lane).

**Figure 5:**
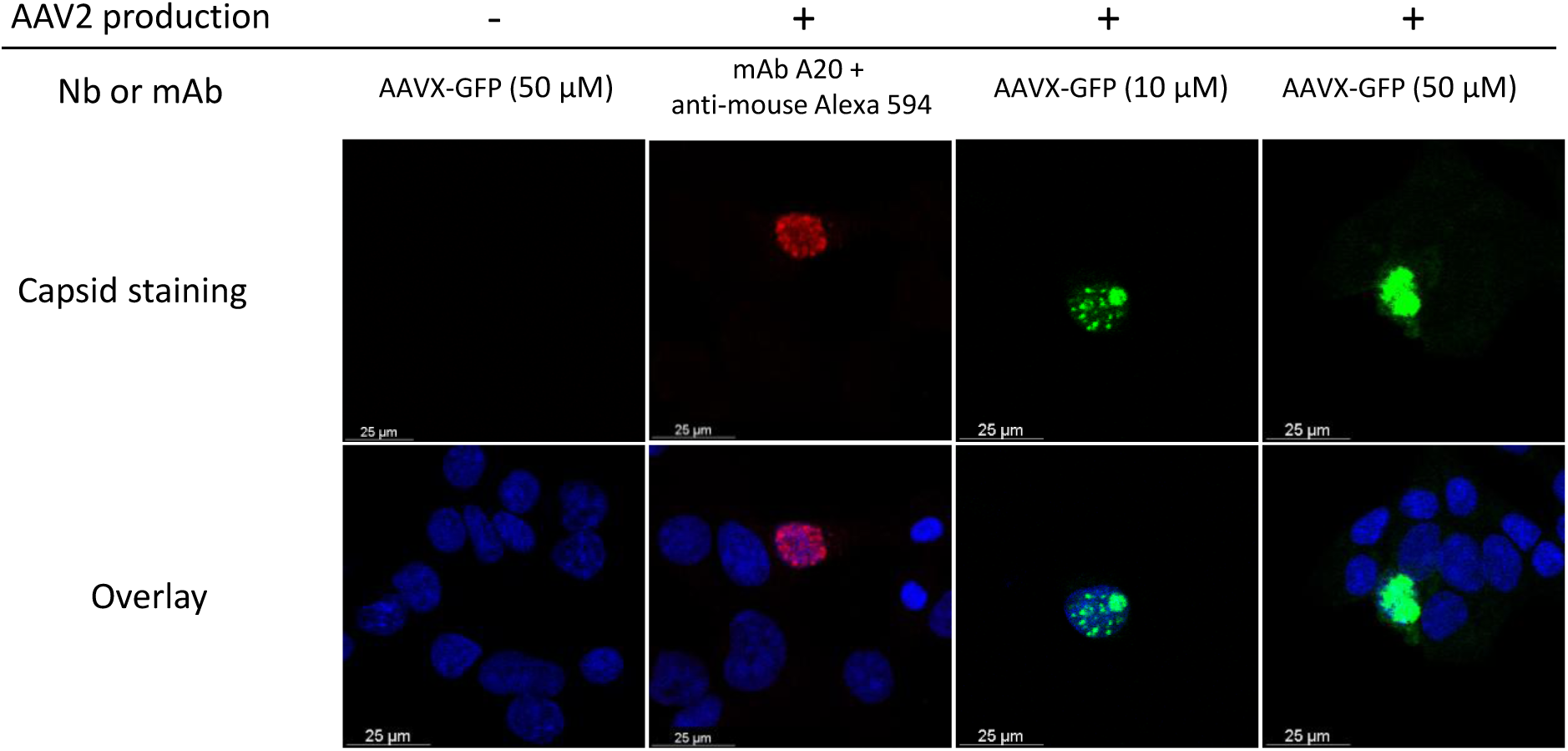
Fluorescence microscopy of HEK-293 cells transfected with pHelper and pRep2Cap2 after fixing and immunostaining with either A20 mAb/ Anti-mouse Alexa Fluor 594 conjugated (red) or AAVX-GFP (green), DAPI staining in blue. 10x magnification.

## Discussion

As our data demonstrate, the Nb AAVX can be produced efficiently as a soluble protein in the cytosol of *E. coli* and can be purified to high purity in a one-step process via IMAC. Cytosolic expression compared to periplasmic expression typically yields higher amounts of nanobody but may increase the risk of protein aggregation. To counteract this, the Origami (DE3) strain was employed to facilitate protein folding through cytoplasmic disulfide bond formation. However, a fraction of the protein was observed in the insoluble fraction, hinting at partial inclusion body formation. The obtained yields after purification are high compared to similar reported productions in shake flasks and one inexpensive, small scale *E. coli* production yields mg amounts of Nb sufficient for hundreds of assays.^35,36^ From the expression yield, one can conclude that Nb AAVX has a very stable folding pathway and final state. Nonetheless, even further improvement possibilities could be employed to refine the production by protein engineering or process optimization.^37–40^ For our research applications, the purity obtained by a single IMAC was sufficient, yet further steps could be added.^41^

Chemical conjugation to a crosslinked agarose resin via lysine residues of the nanobody generated a chromatography material with efficient binding of AAV2 and AAV5 capsids from HEK-293 cell lysates. Titering of the purified capsid samples suggests that the yields are comparable to the commercial AAVX affinity resin. However, we did not test and compare the maximum binding capacities and all concentrations tested were below the manufacturers specified binding capacity of >1×13 vg/mL of resin.^42^ To determine whether a self-made resin is suitable for larger scale or immune critical applications, further analysis of capacity, leaching and endotoxin content should be conducted.

Biolayer interferometry was measured by immobilizing the AAVX-nanobody to streptavidin tips by the fused Strep-tag I and yielded standard curves for the titering of AAV2 and AAV5 capsids. AAV5 capsids exhibited steeper initial association slopes. However, contrary to the AAV2 curves, the concentration-dependent increase remained proportional to AAV5 concentration only during approximately the first 30 seconds of the association phase, after which several sensorgrams intersected. An increased Pluronic F-68 concentration of 0.05% did diminish this effect but did not prevent it (Figure S4), which hints at aggregation as cause of these artifacts. Another noteworthy observation is that affinities could be calculated in a picomolar range, which surpasses the affinity of the Strep-tag I to streptavidin by multiple orders of magnitude, especially when not placed C-terminally.^43^ An explanation for these results may be that sufficient amounts of immobilized nanobody are retained on the surface, despite the low affinity, when switching to the AAV capsids, and that the low affinity to the sensor tip is then mitigated by the multivalency of the capsids.^44^ The overall low apparent dissociation rate also suggests the occurrence of rebinding events enabled by the high ligand concentration and the multivalency of the capsids, as previously reported for other viruses and virus-like particles.^45,46^ Substitution of the Strep-tag I motif for a SBP-tag or an Avi-Tag and biotinylation of the Nb would result in far greater affinity towards the sensor tips and would enable the usage of lower nanobody concentrations that could counteract some of the observed effects for more robust measurements.^47,48^ However, stronger binding might also impede regeneration.

The nanobody proved effective for ELISA either for capturing or detection or both. In case of the nLuc fusion for detection, the reporter signal is not amplified by a secondary binding molecule, yet the efficient enzyme and the sensitive luminescence detection provide a dynamic range comparable to detection with secondary antibodies, while providing the advantage of shorter assay times and eliminating the need for serotype-specific detection reagents for many AAV capsids. Additionally, using AAVX may also serve as a proxy for the affinity of different AAV serotypes toward AAVX-based purification resins.

When comparing BLI and ELISA quantification, the dynamic ranges were in the same ballpark with a narrower tendency for ELISA. However, the saturation point has not been determined for either assay. Quantification with BLI required AAV (analyte) concentrations approximately three orders of magnitude higher than in ELISA, while the required sample volume was much smaller (4 µL for BLI and 100 µL for ELISA). BLI tended to be less time-consuming but required access to special equipment and reagents. Both methods yielded stronger signals for AAV5 than for AAV2 at equivalent concentrations, consistent with previous reports.^29^ Quantification of AAV capsids with AAVX has the advantage that it serves a multitude of serotypes. Additionally, when used with modified capsids and in conjunction with orthogonal quantification, e.g. viral genome titers, the results indicate chromatography behavior for commercial or custom-made AAVX affinity resins.

Immunostaining of AAV particles in HEK-293 cells was achieved using AAVX-GFP as a one-step staining reagent. The fluorescence intensity was comparable to the staining with mAb A20 and the used fluorophore conjugated secondary antibody. Note that monoclonal antibodies are typically used at lower concentrations. The high AAVX-GFP concentrations employed for staining were available due to the efficient production and did not give rise to a significant background signal. Staining with 50 μM AAVX-GFP resulted in diffuse nuclear staining, whereas staining with 10 μM AAVX-GFP or the mAb control showed a spotted fluorescence pattern, which is in line with localization of AAV capsids in nucleoli.^49^ For live cell imaging applications, a DNA plasmid or mRNA with an AAVX expression cassette could be introduced into the cells.^50,51^

The applications for specific research questions can be expanded by chemical conjugation of the AAVX Nb. The protein contains four lysine residues that are not located in or near the CDRs, making lysine-directed conjugation strategies an attractive option. Chemical labelling with fluorophores via lysines may result in multiple labelling with high brightness while retaining a small size, yet hydrophobicity may cause problems.^52^ Introduction of cysteine may provide a more specific handle but can complicate production and purification. Site-specific conjugation can also be achieved by peptide motif incorporation for chemoenzymatic labelling, for example by aldehyde generation through the formylglycine-generating enzyme (FGE) at a cysteine residue within a consensus sequence and subsequent chemical ligation of desired coupling partner.^53,54^ Furthermore, strictly enzymatic approaches can be used e.g. to biotinylate a specific lysine of the AviTag with the *E. coli* biotin ligase (BirA) and open the possibility of using the well-established avidin-biotin interaction.^55^ Depending on the use case, combination tags like SnAvi-Tag could further extend the utility of the nanobody.^56^ Also labeling via protein tags (e.g. SNAP-tag) is an option.^57^

These proof-of-concept experiments show successful utilization of the recombinantly expressed nanobody AAVX for research and bioprocessing applications using two representative serotypes and pave the way for further applications.

## Materials s Methods

### Cloning

Restriction enzymes were purchased from New England Biolabs and T4 Ligase from Thermo Scientific. PCRs for cloning were conducted with Q5 polymerase (NEB) according to the manual. Sequences of all primers used during cloning are listed in table S1 in the supplement. All resulting plasmids were validated by Sanger DNA sequencing. The nucleotide sequence encoding the nanobody AAVX was backtranslated from the amino acid sequence in PDB: 9DC3, optimized for *E. coli* expression, and ordered as a gene from Twist Bioscience. The gene encodes Nb AAVX, a GGSG linker, and a C-terminal His-tag. The sequence that codes for the linker comprises a BamHI recognition site to enable later introduction of fusion protein genes. The gene was inserted into a pET21a(+) backbone by digestion of insert and backbone with NdeI and XhoI restriction enzymes and ligation of both fragments. The final plasmid pET-21a(+)-AAVX-His is referred to as pZMB1127 (Addgene: 254132).

For insertion of eGFP as fusion partner, the eGFP sequence was PCR amplified and equipped with 15 bp homologous regions to the target plasmid with the primers LKR29 and LKR30. The AAVX-His plasmid (pZMB1127) was opened via BamHI and XhoI digest, the backbone purified, and the eGFP fragment was inserted with a HiFi Assembly reaction (NEB). The final plasmid pET-21a(+)-AAVX-eGFP-His is referred to as pZMB1128 (Addgene: 254133).

The dual His-tag was added by PCR amplification of the Nb AAVX sequence from the AAVX-His plasmid (pZMB1127) with the overhang primers SEQ250 and LKR36. The amplified gene and the AAVX-GFP plasmid were restricted with NdeI and XhoI, purified, and ligated yielding a plasmid that contains two Hexahistidine-tags that are linked by a Strep-tag I motif C-terminal of the AAVX Nb. The final plasmid pET-21a(+)-AAVX-His-StrepI-His is referred to as pZMB1133 (Addgene: 254134).

To insert the nLuc as fusion partner, the DNA sequence was taken from pNL1.1 (Promega) and the gene was ordered as gene synthesis (GeneArt/Thermo Fisher Scientific) and PCR amplified with overhang primers (LKR34 and LKR35) containing a BamHI and a XhoI restriction site. The correspondingly digested PCR product and AAVX-GFP plasmid were purified and ligated. The final plasmid pET-21a(+)-AAVX-nLuc-His is referred to as pZMB1134 (Addgene: 254135).

### Protein expression and purification

Chemicals used were purchased in p.a. grade from Carl Roth unless stated otherwise. All Nb AAVX plasmid constructs were transformed into *E. coli* Origami (DE3) and single colonies were grown in 500 mL LB-medium (LB-Broth Lennox, Roth) supplemented with 100 µg/mL ampicillin in baffled 2 L Erlenmeyer flasks with orbital shaking at 180 rpm and 24 °C until OD_600_ reached 0.8. Protein expression was induced by adding IPTG (isopropyl β-D-thiogalactopyranosid) to a final concentration of 1 mM and cultures were incubated for another 18 h at 24 °C. Cells were harvested by centrifugation at 8000×g, 4°C for 3 min and pellets were resuspended in chilled lysis buffer (50 mL/L culture; 50 mM Na_2_HPO_4_, 300 mM NaCl, 2.5 mM MgCl_2_, 0.5 mM CaCl_2_, 1 mM PMSF (Sigma Aldrich), 1 U/mL DNaseI, pH 7.4). The cell membranes were disrupted by three cycles in a French press (SLM Aminco FA-078 with a FA-032 cell) at a pressure of 144 MPa (approx. 20800 psi) and cell debris and insoluble protein were separated from the soluble fraction by centrifugation at 10000×g, 4°C for 10 min and a second centrifugation step at 40000×g, 4°C for 30 min. After filtration (0.45 µm PES syringe filter, Sartorius), the supernatant was applied to a Ni-NTA agarose (Macherey Nagel) column (1.5 mL resin, 10 mm diameter) that was in equilibration buffer (50 mM Na_2_HPO_4_, 300 mM NaCl, pH 7.4) and connected to an Äkta Start chromatography system (Cytiva). The column was washed with 5 column volumes (CV) equilibration buffer followed by 15 CV equilibration buffer supplemented with 20 mM imidazole. Protein was eluted with high-imidazole buffer (50 mM Na_2_HPO_4_, 300 mM NaCl, 300 mM imidazole, pH 7.4). Protein constructs with dual His-tags were eluted with a similar buffer that contained 1 M imidazole instead of 300 mM. Eluate fractions that contained protein based on 280 nm absorption measurement, were pooled and buffer was exchanged either by dialysis against a 1000-fold volume of PBS (137 mM NaCl, 2.6 mM KCl, 10 mM Na_2_HPO_4_, 1.8 mM KH_2_PO_4_, pH 7.4) overnight at 4°C in a dialysis membrane (6-8 kDa MWCO, Spectra/Por, Repligen) or with an Amicon Ultracentrifugal filter (10 kDa MWCO, Merck Millipore) if the POI had a molecular weight greater than 30 kDa. Final protein concentrations were determined via spectrophotometry (NanoDrop, Thermo Fisher) based on calculated extinction coefficients at 280 nm (for extinction coefficients see supplement table S4).

### AAV production

15 cm cell culture dishes were seeded with 9×10^6^ HEK-293 cells (DSMZ - German Collection of Microorganisms and Cell Cultures) in 20 mL DMEM (Sigma Aldrich) + 10% (v/v) FCS (Sigma Aldrich) + 1% (v/v) Penicillin/Streptomycin (Sigma Aldrich) and incubated at 37°C, 5% CO_2_ 24 h before transfection. For viral particle production, cells were triple-transfected *via* calcium phosphate with a total amount of 45 μg DNA consisting of pHelper (pZMB0088), pGOI containing a CMV-GFP-hGHpA expression cassette flanked by ITRs (pZMB0990), and the pRep2Cap2 (pZMB0916) or pRep2Cap5 (pZMB0647), respectively, in a molar 1:1:1 ratio. ^58–60^

### AAV affinity purification

Cells were harvested 72 h post transfection by scraping from the plates and centrifugation at 3000×g for 5 min. Pelleted cells were resuspended in 0.5 mL lysis buffer (50 mM Tris, 150 mM NaCl, 2 mM MgCl2, pH 7.5) and lysed in three freeze-thaw cycles (−80°C freezer, 15 min /37°C thermo mixer, 5 min). The cell lysate was treated with 0.2 µL Denarase (c-Lecta) to a final concentration of 100 U/mL for 30 min at 37°C, followed by addition of 25 µL CHAPS (3-[(3-cholamidopropyl)dimethylammonio]-1-propane sulfonate, 0.5% w/v final) and further 30 min incubation at 37°C. Lysate was cleared by centrifugation at 3000×g, 4°C for 10 min and the supernatant was saved and stored at -80°C until further use. AAV particles were purified using POROS CaptureSelect AAVX affinity resin (Thermo Fisher Scientific) in a batch procedure with 100 µL resin for each 15 cm plate in 5 mL reaction tubes. AAVX-POROS resin was equilibrated two times with 10 CV PBS, followed by incubation with cell lysate supernatant for 30 min on a tube roller mixer. After each step the resin was pelleted by centrifugation at 4000×g for 2 min to separate it from the supernatant. The resin was washed with 10 CV PBS-T (PBS+0.05% Tween 20) and bound particles were eluted with 2 CV 0.1 M glycine-HCl (pH 2.2) for 10 min with an immediate neutralization of the eluate fraction with 2 M Tris (Fisher Scientific). The final AAV solution was concentrated and rebuffered with an Amicon Ultracentrifugal filter (100 kDa MWCO, Merck Millipore) to HBSS (Sigma Aldrich) and stored at -80°C until use.

### AAV genomic titer determination via qPCR

The viral titer was determined by real-time qPCR. For this, 5 μL AAV solution were treated with 10 U DNaseI (NEB) in DNaseI reaction buffer (NEB) in a reaction volume of 50 μL for 30 min at 37°C, followed by DNase inactivation at 75°C for 20 min, and dilution with 200 μL nuclease-free water supplemented with 0.05% (w/v) Pluronic F68. 2.5 μL samples were mixed with 1.25 µL primer qPCR-CMV-rev (5’-GGCGGAGTTGTTACGACA-3’) and 1.25 µL primer qPCR-CMV-for (5’-GGGACTTTCCTACTTGGCA-3’) with a final primer concentration of 500 nM each, and 5 μL 2× GoTaq qPCR Mastermix (Promega). The qPCR was performed in a real-time PCR instrument (CFX Opus 96, Bio-Rad) with the settings: 10 min at 95°C, 40 cycles with 15 s at 95°C and 30 s at 60°C. The genomic titer was calculated from an internal standard curve that was created from a serial dilution of the pGOI plasmid.

### AAVX covalent immobilization

AAVX-GFP was covalently coupled to AminoLink resin (Thermo Scientific) by lysine-aldehyde reaction. For this, 0.5 mL resin were transferred to a 5 mL reaction tube and equilibrated with 3 CV PBS (pH: 7.4) on a tube roller mixer for 60 min at 4°C. After every incubation step, the resin was centrifuged at 4000×g, 2 min, 4°C and the supernatant was sampled for protein content and discarded. 5 mg AAVX-GFP were diluted in 750 µL PBS and 100 mM NaCNBH_3_ (Sigma Aldrich) and incubated at 4°C on the tube roller overnight. The resin slurry was washed with 2 CV PBS and subsequently quenched by addition of 1 CV of 1 M Tris with 50 mM NaCNBH_3_ and incubation at RT for 30 min on the tube roller. The resin was washed with 5 CV 1 M NaCl, followed by 3 CV PBS.

### AAV capsid sandwich ELISA with AAVX-nLuc detection

For the detection of intact capsids via ELISA, white 96-well high binding ELISA plates (Sarstedt) were coated in 100 μL bicarbonate buffer (pH 9.6) overnight at 4°C with either 1 µg AAVX-His or PKD1-MBP or PKD2-MBP for capturing AAV5 or AAV2, respectively. The following day, the wells were washed 3 times (200 μL PBS + 0.05% Tween20) and blocked with 0.8% BSA (Sigma Aldrich) in PBS for 1 h at RT and washed (3× 200 μL PBS-T). AAV samples were diluted in PBS to a final volume of 100 μL and incubated in the wells for 1 h at RT. After three washing steps (200 μL PBS-T), 250 ng AAVX-nLuc diluted in 100 μL PBS was added to each well and incubated at RT for 1 h. The wells were washed four times with 200 μL PBS-T. Nano-Glo Substrate (Promega) was prepared in Assay Buffer according to the manufacturer’s description, diluted 1:10 in PBS, and 100 μL were added to each well. After 10 min settling time, the luminescence signal was recorded with a plate reader (Infinite M Plex, Tecan).

### AAV capsid sandwich ELISA with Nb AAVX coating and mAb detection

For quantification of AAV capsids in a sandwich ELISA with AAVX-His as capturing nanobody, a clear 96-Well plate (Nunc MaxiSorb, Thermo Scientific) was coated with varying AAVX-His concentrations in 100 μL bicarbonate buffer (pH 9.6) and incubated overnight at 4°C. The following day, the wells were washed (3× 200 μL PBS-T) and blocked with 0.8% BSA in PBS for 1 h at RT. After washing (3× 200 μL PBS-T), AAV samples diluted in 100 µl PBS were added and incubated for 1 h at RT. The washing steps were repeated and A20 antibody (#690055, Progen) to detect intact AAV2 capsids or ADK5a antibody (Progen) to detect intact AAV5 capsids diluted in 100 μL PBS + 0.8% BSA were added to the wells for 1 h at RT. After discarding the primary antibody and washing the wells (3× 200 μL PBS-T), an anti-mouse HRP-conjugated antibody (#31450, ThermoFisher Scientific) diluted in 100 μL PBS + 0.8% BSA was added to the wells and incubated for 1 h at RT. Subsequently, unbound secondary antibody was discarded and wells were washed four times with 200 μL PBS-T. Following that, ABTS buffer (3.25 mM BH_2_NaO_4_, 40 mM citric acid, 60 mM Na_2_HPO_4_, pH 4.5) with 1 g/L ABTS (2,2’-azino-bis[3-ethylbenzothiazoline-6-sulfonic acid], Biochemica) was added and after 30 min absorbance at 405 nm was measured in a plate reader (Infinite M Plex, Tecan).

### Biolayer Interferometry Measurements

Samples were analyzed on a BLItz N1 (ForteBio; now Sartorius) system with Octet Streptavidin Biosensors (Sartorius). Sensor tips were hydrated for 10 min in assay buffer prior to a measurement at 21°C and AAVX-His-StrepI-His samples and AAV2 or AAV5 were freshly diluted in assay buffer (10 mM Na_2_HPO_4_, 1.8 mM KH_2_PO_4_, 2.6 mM KCl, 200 mM NaCl, 0.1% BSA, 0.001% Pluronic F68, pH 7.4). The following instrument settings were used: 60 s initial baseline step in assay buffer, 300 s loading step with 1 μM AAVX-dHis, 120 s baseline step in assay buffer, 300 s association step, 300 s dissociation step in assay buffer, 120 s regeneration step in regeneration buffer (100 mM glycine-HCl, pH 2.2) and 60 s baseline step in assay buffer. Data analysis for calculation of the kinetic constants was performed using the Octet N1 Software v 1.4.0.13. For the generation of a concentration calibration curve, the slopes of the initial linear phases of the association steps were plotted against the concentration and fitted with a four-parameter logistic (4PL) regression using the software Origin Pro 2018.

### Immunostaining

8-Well chamber Slides (Ibidi) were rinsed, dried and coated with 200 μL poly-D-lysine (Sigma Aldrich). 1.5×10^4^ HEK-293 cells in 200 μL DMEM + 10% (v/v) FCS + 1% Penicillin/Streptomycin were seeded in each well and incubated at 37°C, 5% CO_2_. After 24 h, cells were transfected with pRep2Cap2 and pHelper *via* calcium phosphate transfection with a total of 85 ng DNA/well. 48 h post transfection, the cells were washed twice with PBS, fixed with 4% paraformaldehyde (PFA) at RT for 5 min, washed and permeabilized with 0.1% Triton X-100 (Sigma Aldrich) in H_2_O for 5 min at RT. The cells were washed again (2× 200 μL PBS) and subsequently blocked with 200 μL 3% BSA in PBS for 30 min at RT and the blocking solution was discarded. For staining, 10 μM or 50 µM AAVX-GFP solution were added to the cells and incubated overnight at 4°C. As staining control, the procedure was performed with monoclonal antibody A20 (Progen). On the next morning, only in the case of the antibody A20, solution was discarded, cells washed with PBS, and 200 μL anti-mouse Alexa Fluor 594 conjugated antibody (Thermo Fisher Scientific) were added for 1 h at RT. For both, AAV-GFP and A20 samples, wells were washed (2× 200 μL PBS) and cells counterstained with 10 μM DAPI (AppliChem) in PBS for 10 min at RT. Finally, the cells were washed (2× 200 μL PBS) and stored in PBS in the dark until fluorescence confocal microscopy imaging (Stellaris 8 Falcon; Leica).

### Protein thermal stability assay

Protein solutions were diluted in PBS (pH 8.0) and mixed with ProteOrange (LumiProbe) to a final dye concentration of 5× in a total sample volume of 25 µL. All samples were transferred to a white 96-well qPCR microtiter plate and sealed with an adhesive film. Fluorescence was recorded using a Real-Time PCR system (CFX Opus 96, Bio-Rad). Thermal unfolding was monitored during heating from 25°C to 95°C in 0.5°C increments with 30 seconds incubation for each temperature step.

## Data availability statement

Data are provided with this paper. Source data are available upon request from the corresponding author. All plasmids generated in this study have been deposited with Addgene using Plasmid IDs 254132-254135.

## Acknowledgments

We acknowledge support for the publication costs by the Open Access Publication Fund of Bielefeld University. We thank Philipp Borchert for technical support.

## Author contributions

L.H.: Conceptualization, Investigation, Methodology, Validation, Visualization, Writing – original draft, Writing – review & editing; M.F.G: Investigation; K.M.M: Conceptualization, Methodology, Validation, Writing – review C editing, Supervision

