## Supplementary Information for "*E. coli*-produced nanobody fusions as a pan-serotype platform for AAV purification, quantification, and detection"

*Table S1: Nucleic acid sequences of all oligonucleotides used for cloning and qPCR.*

| **Name** | **Sequence** |
|---|---|
| **LKR29 eGFP Gibson for** | GTTACCGTATCCAGCGGCGGATCCGGCGTGAGCAAGGGCGAGGAGC |
| **LKR30 eGFP Gibson rev** | GTGGTGGTGGTGGTGGTGCTCGAGACCCTTGTACAGCTCGTCCATG |
| **LKR34 nLuc-BamHI for** | AAAAAGGATCCGGCGCCCCAGTCTTCACACTCG |
| **LKR35 nLuc XhoI rev** | AAAAACTCGAGGCCTCCCGCCAGAATGCGTTCGC |
| **LKR36 dHis6 XhoI rev** | TTTTTCTCGAGACCGCCAAACTGGGGATGACGCCAGGCCCGTGAGTGAT<br>GATGATGATGATGACCGGAGCCGCCGCTGGATACGGTAACCTGCGTACC |
| **SEQ250 dHis6 for** | CCACGATGCGTCCGGCGTAG |
| **qPCR CMV for** | GGGACTTTCCTACTTGGCA |
| **qPCR CMV rev** | GGCGGAGTTGTTACGACA |

*Table S2: Nucleic acid sequences of the Nb AAVX and the fused domains. Domains and motifs are color coded: Nb AAVX: blue, GS-linker: grey, His-tag: orange,Strep tag I: purple, NLuc: yellow, EGFP: green.*

| **Domain** | **Nucleic acid sequence** |
|---|---|
| **AAVX** | ATGCAGGTACAGATCCAGGAAAGCGGAGGCGGGATCGTTCAGGCCGGCGGCTCACTTCGGCTGTCCTGCGCGGC<br>CAGCGGGCGCACCCATGGTATGTATGCGATGGGTTGGTTCAGACAGGCACCGGGTAAAGAACGAGAATTTGTTGCA<br>GTGCAGGACATCACAGCCTCAAATACACACTACAGCTCGGCTGTGAAGGGTCGGTTCACTCTTTCAAGAGACAATGC<br>AAAAAATACAGCCTACCTCCAAATGAATAACCTGAAACCTGAAGATACGGCAGTGTATTATTGCGCTGCTGGACCTAC<br>ATTAATGTCAGGTTCGTACAACTCCGCCCGCGACTATGATTACTGGGGTCAAGGTACGCAGGTTACCGTATCCAGC |
| **-His** | GGCGGATCCGGCCATCACCATCATCAT |
| **-dHis** | GGCGGCTCCGGTCATCATCATCATCATCACTCACGGGCCTGGCGTCATCCCCAGTTTGGCGGTCTCGAGCACCAC<br>CACCACCACCAC |
| **-NLuc** | GGCGGATCCGGCGCCCCAGTCTTCACACTCGAAGATTTCGTTGGGGACTGGCGACAGACAGCCGGCTACAACCTG<br>GACCAAGTCCTTGAACAGGGAGGTGTGTCCAGTTTGTTTCAGAATCTCGGGGTGTCCGTAACTCCGATCCAAAGGATT<br>GTCCTGAGCGGTGAAAATGGGCTGAAGATCGACATCCATGTCATCATCCCGTATGAAGGTCTGAGCGGCGACCAAAT<br>GGGCCAGATCGAAAAAATTTTTAAGGTGGTGTACCCTGTGGATGATCATCACTTTAAGGTGATCCTGCACTATGGCACA<br>CTGGTAATCGACGGGGTTACGCCGAACATGATCGACTATTTCGGACGGCCGTATGAAGGCATCGCCGTGTTCGACG<br>GCAAAAAGATCACTGTAACAGGGACCCTGTGGAACGGCAACAAAATTATCGACGAGCGCCTGATCAACCCCGACGG<br>CTCCCTGCTGTTCCGAGTAACCATCAACGGAGTGACCGGCTGGCGGCTGTGCGAACGCATTCTGGCGGGAGGCCT<br>CGAGCACCACCACCACCACCAC |
| **-GFP** | GGCGGATCCGGCGTGAGCAAGGGCGAGGAGCTGTTCACCGGGGTGGTGCCCATCCTGGTCGAGCTGGACGGCG<br>ACGTAAACGGCCACAAGTTCAGCGTGTCCGGCGAGGGCGAGGGCGATGCCACCTACGGCAAGCTGACCCTGAAG<br>TTCATCTGCACCACCGGCAAGCTGCCCGTGCCCTGGCCCACCCTCGTGACCACCCTGACCTACGGCGTGCAGTG<br>CTTCAGCCGCTACCCCGACCACATGAAGCAGCACGACTTCTTCAAGTCCGCCATGCCCGAAGGCTACGTCCAGGA<br>GCGCACCATCTTCTTCAAGGACGACGGCAACTACAAGACCCGCGCCGAGGTGAAGTTCGAGGGCGACACCCTGGT<br>GAACCGCATCGAGCTGAAGGGCATCGACTTCAAGGAGGACGGCAACATCCTGGGGCACAAGCTGGAGTACAACTA<br>CAACAGCCACAACGTCTATATCATGGCCGACAAGCAGAAGAACGGCATCAAGGTGAACTTCAAGATCCGCCACAAC<br>ATCGAGGACGGCAGCGTGCAGCTCGCCGACCACTACCAGCAGAACACCCCCATCGGCGACGGCCCCGTGCTGC<br>TGCCCGACAACCACTACCTGAGCACCCAGTCCGCCCTGAGCAAAGACCCCAACGAGAAGCGCGATCACATGGTC<br>CTGCTGGAGTTCGTGACCGCCGCCGGGATCACTCTCGGCATGGACGAGCTGTACAAGGGTCTCGAGCACCACCAC<br>CACCACCAC |

*Table S3: Amino acid sequences of the Nb AAVX and the fused domains. Domains and motifs are color coded: Nb AAVX: blue, GS-linker: grey, His-tag: orange,Strep tag I: purple, NLuc: yellow, EGFP: green.*

| Domain | Amino acid sequence |
|---|---|
| AAVX | MQVQIQESGGGIVQAGGSLRLSCAASGRTHGMYAMGWFRQAPGKEREFVAVQDITASNTHYSSAVKGRFTLSRDNAKNTA YLQMNNLKPEDTAVYYCAAGPTLMSGSYNSARDYDYWGQGTQVTVSS |
| -His | GGSGHHHHH |
| -dHis | GGSGHHHHHHSRAWRHPQFGGLEHHHHHH |
| -NLuc | GGSGAPVFTLEDFVGDWRQTAGYNLDQVLEQGGVSSLFQNLGVSVTPIQRIVLSGENGLKIDIHVIIPYEGLSGDQMGQIEKIF KVVYPVDDHHFKVILHYGTLVIDGVTPNMIDYFGRPYEGIAVFDGKKITVTGTLWNGNKIIDERLINPDGSLLFRVTINGVTGWR LCERILAGGLEHHHHHH |
| -GFP | GGSGVSKGEELFTGVVPILVELDGDVNGHKFSVSGEGEGDATYGKLTLKFICTTGKLPVPWPTLVTTLTYGVQCFSRYPDHMK QHDFFKSAMPEGYVQERTIFFKDDGNYKTRAEVKFEGDTLVNRIELKGIDFKEDGNILGHKLEYNYNSHNVYIMADKQKNGIK VNFKIRHNIEDGSVQLADHYQQNTPIGDGPVLLPDNHYLSTQSALSKDPNEKRDHMVLLEFVTAAGITLGMDELYKGLEHHH HHH |

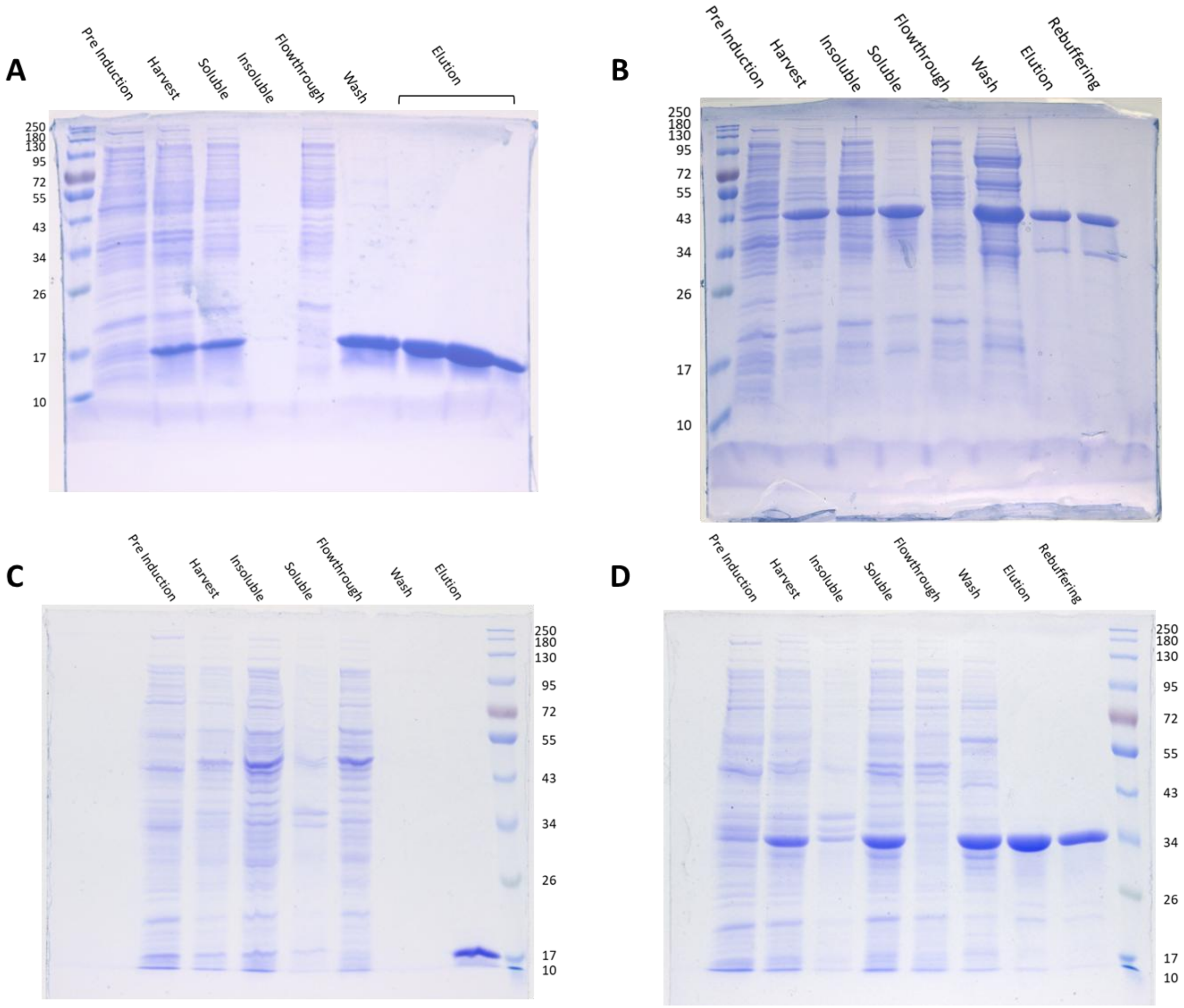


*Figure S1: SDS-gels characterizing productions and purifications of Nb AAVX and fusion variants. (A) AAVX-His (B) AAVX-GFP (C) AAVX-dHis (D) AAVX-NLuc; Pre Induction: Whole cell protein before induction; Harvest: Whole cell protein 18 h after induction; Insoluble: pelleted cell debris after cell lysis; Soluble: soluble proteins after cell lysis; Flowthrough: flowthrough of cell lysate during IMAC; Wash: wash fraction during IMAC; Elution: pooled elution fractions; Rebuffering: purified protein after rebuffering and concentration.*

*Table S4: Protein masses and extinction coefficients calculated based on the Pace method and used for molar and mass concentration determination.*

| Protein | Mass / Da | Extinction coefficient ε / L·mol$^{-1}$·cm$^{-1}$ |
|---|---|---|
| **AAVX-His** | 14692 | 23045 |
| **AAVX-GFP** | 41921 | 45060 |
| **AAVX-dHis** | 17175 | 28545 |
| **AAVX-NLuc** | 34300 | 48485 |

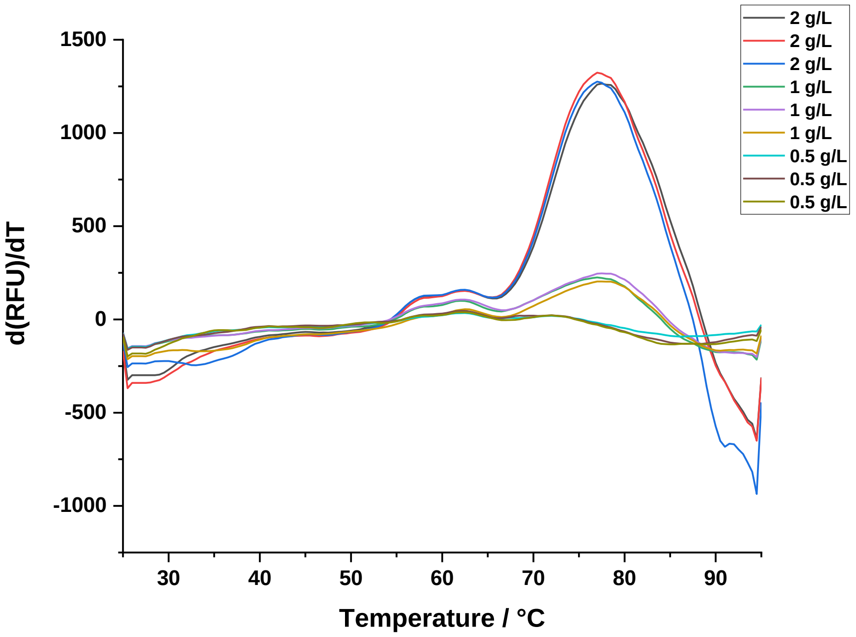


*Figure S2: First derivative of the ProteOrange fluorescence for AAVX melting curve with concentrations from 0.5 to 2 g/L.*

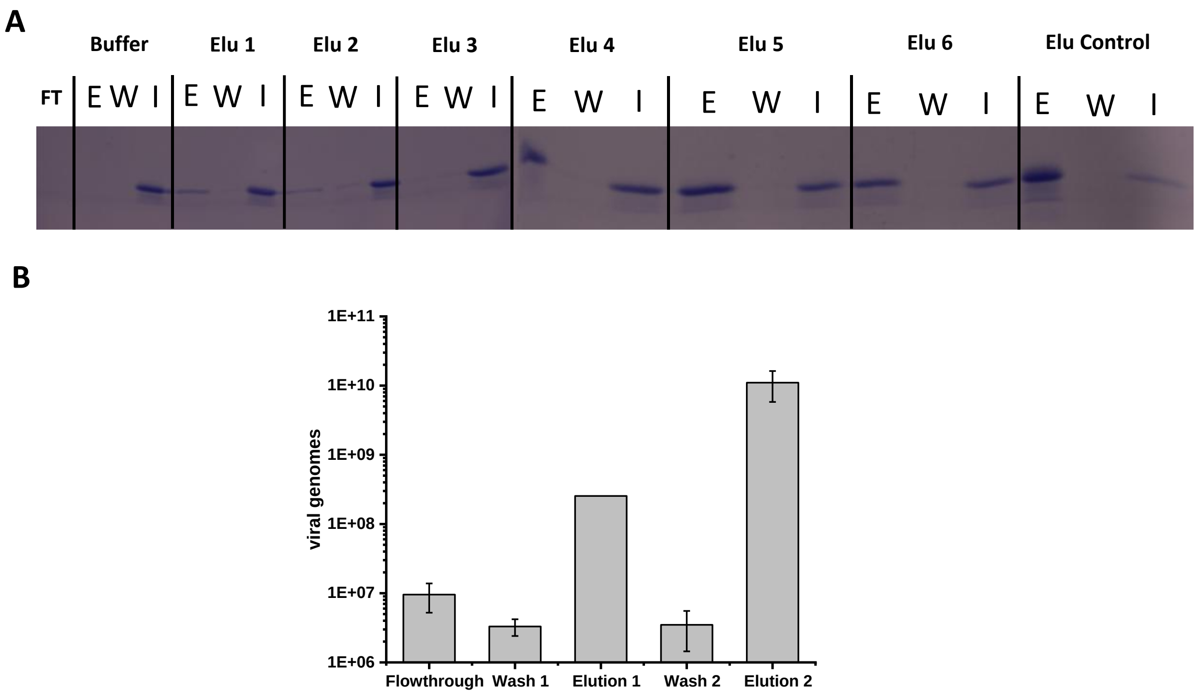


*Figure S3: (A)SDS-Gel with supernatant sample from tested elution conditions (Table S5) to check if AAVX is eluted from Ni-NTA resin; E: Tested elution condition; W: Wash step; I: elution step with imidazole buffer. (B) qPCR of elution process with buffer Elu 3.*

*Table S5: Buffer compositions that were tested in the elution steps for Ni NTA purification attempts.*

| Name | Buffer | Salt | pH |
|---|---|---|---|
| **Buffer** | 50 mM $Na_2HPO_4$ | 300 mM NaCl | 7.4 |
| **Elu 1** | 100 mM NaAc | 1.5 M $MgCl_2$ | 5 |
| **Elu 2** | 100 mM NaAc | 2.5 M $MgCl_2$ | 4 |
| **Elu 3** | 100 mM NaAc | 0.5 M $MgCl_2$ | 4 |
| **Elu 4** | 100 mM NaAc | 2.5 M $MgCl_2$ | 3 |
| **Elu 5** | 100 mM NaAc | 1.5 M $MgCl_2$ | 3 |
| **Elu 6** | 100 mM NaAc | 0.5 M $MgCl_2$ | 3 |
| **Elu Control** | 100 mM Glycin-HCl | - | 2.2 |

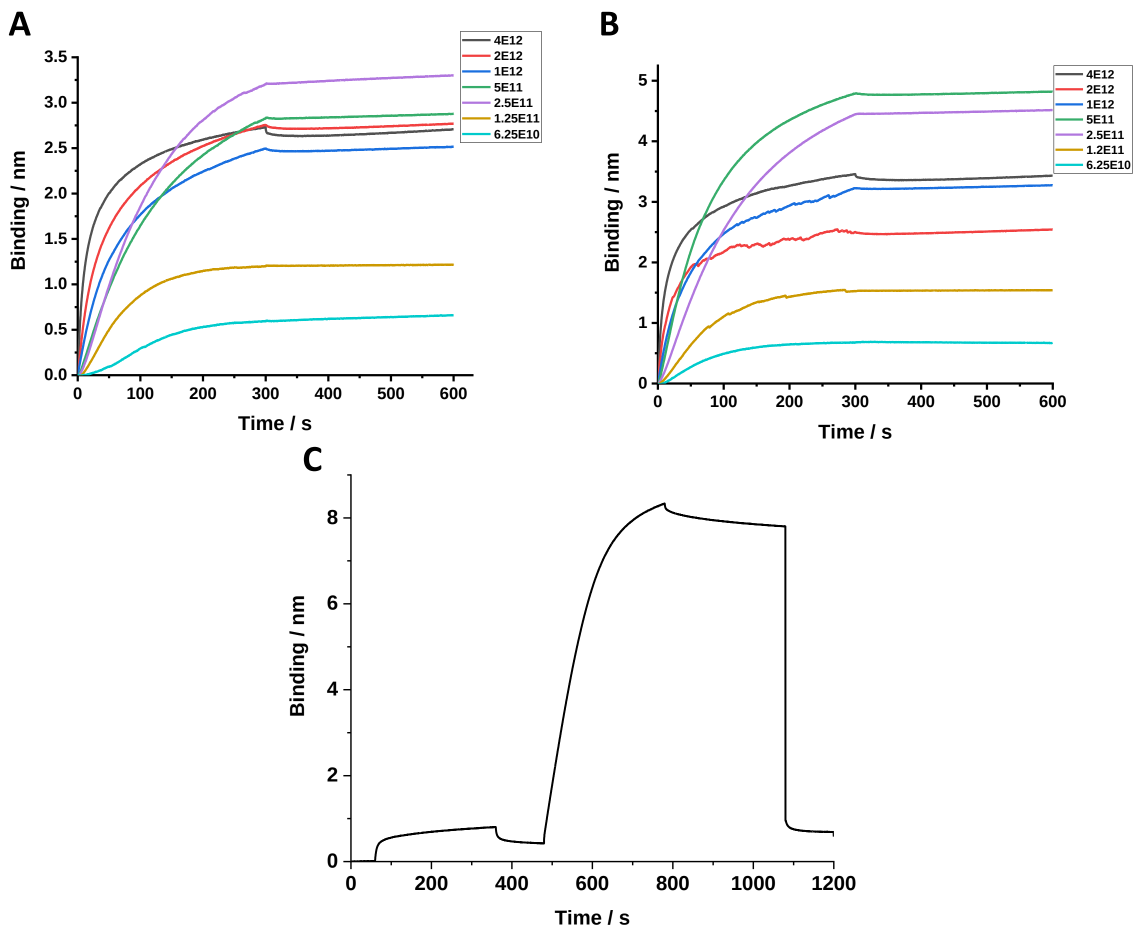


*Figure S4: BLI Sensorgrams of association and dissociation of the Nb AAVX and AAV5 capsid interactions with concentrations ranging from 6.25×10^10 to 4×10^12 vg/mL. With the BLI buffer containing. (A) 0.05 % Pluronic F-68 or (B) 0.001% Pluronic F-68. (C) Full sensorgram with the steps AAVX-dHis loading on streptavidin sensor, wash, AAV5 association, dissociation, and regeneration.*

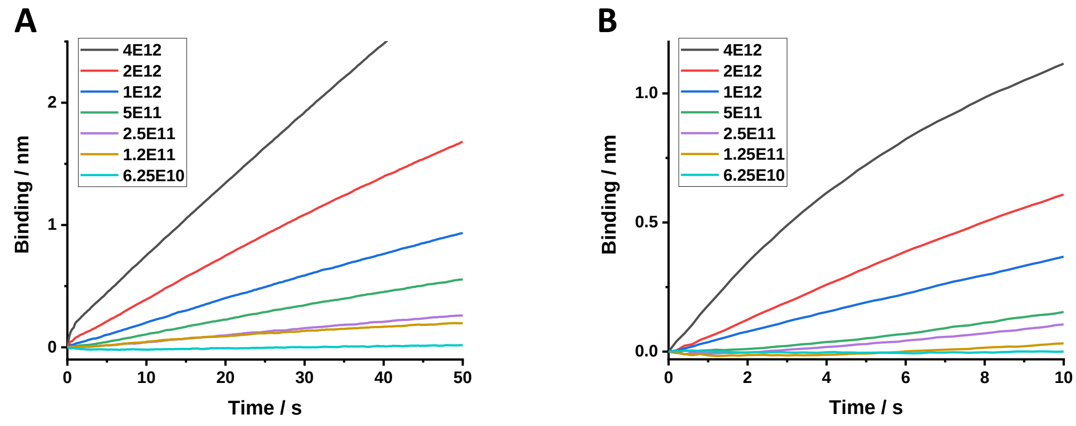


*Figure S5: Sensorgrams of the initial association step of varying particle concentrations ranging from 6.25×10^10 to 4×10^12 vg/mL of (A) AAV2 (B) AAV5.*
